# Multi-session transcutaneous spinal cord stimulation prevents chloride homeostasis imbalance and the development of spasticity after spinal cord injury in rat

**DOI:** 10.1101/2023.10.24.563419

**Authors:** Dillon C. Malloy, Marie-Pascale Côté

## Abstract

Spasticity is a complex and multidimensional disorder that impacts nearly 75% of individuals with spinal cord injury (SCI) and currently lacks adequate treatment options. This sensorimotor condition is burdensome as hyperexcitability of reflex pathways result in exacerbated reflex responses, co-contractions of antagonistic muscles, and involuntary movements. Transcutaneous spinal cord stimulation (tSCS) has become a popular tool in the human SCI research field. The likeliness for this intervention to be successful as a noninvasive anti-spastic therapy after SCI is suggested by a mild and transitory improvement in spastic symptoms following a single stimulation session, but it remains to be determined if repeated tSCS over the course of weeks can produce more profound effects. Despite its popularity, the neuroplasticity induced by tSCS also remains widely unexplored, particularly due to the lack of suitable animal models to investigate this intervention. Thus, the basis of this work was to use tSCS over multiple sessions (multi-session tSCS) in a rat model to target spasticity after SCI and identify the long-term physiological improvements and anatomical neuroplasticity occurring in the spinal cord. Here, we show that multi-session tSCS in rats with an incomplete (severe T9 contusion) SCI (1) decreases hyperreflexia, (2) increases the low frequency-dependent modulation of the H-reflex, (3) prevents potassium-chloride cotransporter isoform 2 (KCC2) membrane downregulation in lumbar motoneurons, and (4) generally augments motor output, i.e., EMG amplitude in response to single pulses of tSCS, particularly in extensor muscles. Together, this work displays that multi-session tSCS can target and diminish spasticity after SCI as an alternative to pharmacological interventions and begins to highlight the underlying neuroplasticity contributing to its success in improving functional recovery.

## INTRODUCTION

There are extensive secondary complications that follow a spinal cord injury (SCI), and the development, persistence, and severity of such conditions can be widely heterogeneous throughout the SCI population. Many of these complications are instigated by dysfunction in the sensorimotor system as the integration between sensory and motor components is disrupted. Amongst those is the development of spasticity that affects nearly 75% of the SCI population (Holtz et al., 2017; Maynard et al., 1990). Consequences of spasticity include both an increase in spinal excitation and impaired spinal inhibition, which both contribute to the development of spastic symptoms such as involuntary muscle movements, co-contractions of antagonistic muscles, hyperactive reflexes, and the inability to depress stretch reflexes in a context-dependent manner (Bennett et al., 1999; Malmsten, 1983; Nielsen et al., 2007; Valero-Cabre et al., 2004). Standard practice to treat spasticity after SCI is through pharmacological intervention with anti-spastic medications that mostly act centrally to depress the net neural output, thereby decreasing reflex hyperexcitability at the cost of reduced muscle activity and muscle weakness (Dietz et al., 2023). This is a hampering approach that must be improved; therefore, it is integral that new therapeutic strategies target the sensorimotor system and restore spinal reflex inhibition without negatively impacting functional motor excitability and output.

Emerging popularity of therapeutic strategies like neuromodulation of the spinal cord with electrical stimulation through transcutaneous spinal cord stimulation (tSCS) after injury has become increasingly studied with reports suggesting a potential for meaningful functional recovery for SCI individuals. The focal point of these interventions has largely been geared toward restoring volitional movements in the legs (Gerasimenko et al., 2015a; 2015c; Sayenko et al., 2019; Taccola et al., 2018) and arms (Gad et al., 2018; Inanici et al., 2018; 2021) as well as improving locomotion (Gad et al., 2017; Hofstoetter et al., 2015; Minassian et al., 2016a). However, recent studies suggest that single tSCS sessions also have the potential to decrease spasticity (Estes et al., 2017; Hofstoetter et al., 2014; 2020; Sandler et al., 2021). While these early studies suggest a potential for tSCS to decrease spasticity after SCI, there is a need to refocus outcome measures on spasticity alone and to more precisely identify the specific pathways and mechanisms involved. Also, it will be critical to understand the potential effects of repeated and sustained sessions of tSCS, rather than a single stimulation session, to determine how this intervention can be used to treat or diminish spasticity after SCI. In addition, while human-based investigations provide critical information on functional recovery, they are limited in only providing indirect evidence on the neurophysiological mechanisms and do not offer insights into the neuroanatomical plasticity induced by tSCS that contribute to decrease spasticity and spastic-like symptoms after SCI. The advantage of using an animal model is the ability to directly identify the spinal pathways and molecular mechanisms involved with using this intervention.

An imbalance in chloride homeostasis following spinal cord injury has been well established to contribute to reflex disinhibition, reflex hyperexcitability, and the development and maintenance of spasticity (Boulenguez et al., 2010; Côté et al., 2014; Gackiere and Vinay, 2015). The membrane-bound protein responsible for chloride extrusion, potassium-chloride cotransporter isoform 2 (KCC2), has been shown to be internalized and downregulated on the motoneuronal membrane following SCI, which results in disrupted chloride homeostasis and increased excitability of the reversal potential for inhibitory postsynaptic potentials (Boulenguez et al., 2010). This can be mitigated by activity-based therapy after SCI as exercise can restore chloride homeostasis by returning motoneuronal KCC2 membrane expression which, in turn, contributes to decreased reflex hyperexcitability and improved reflex modulation (Beverungen et al., 2020; Bilchak et al., 2021; Côté et al., 2014), all of which are important mechanisms contributing to spasticity. tSCS, like exercise, functions to repetitively activate primary afferents, so it can be used as an alternative activity-based therapy to promote activity-dependent plasticity. This also suggests that tSCS may share common neuroanatomical mechanisms with activity-based therapy. Since restoring motoneuronal chloride homeostasis has been shown to be integral for mitigating reflex hyperexcitability with activity-based therapy (Beverungen et al., 2020; Bilchak et al., 2021; Côté et al., 2014), it highlights the importance of investigating the impact of tSCS to alter motoneuronal chloride homeostasis after SCI, particularly KCC2.

Previous work from our laboratory has established and validated the feasibility of successfully delivering tSCS over multiple sessions in awake rats with a complete spinal cord transection and showed that it induces an increase in motor output in ankle muscles (Malloy et al., 2022), similar to human SCI individuals (Murray and Knikou, 2019). To bolster the clinical relevance of our model, we progressed our injury type from a complete transection to a contusion injury to focus on anatomically incomplete SCI, and we expanded our investigations into the effect of tSCS on spinal hyperreflexia and its contribution to improving spasticity-related outcome measures after SCI. Here, we show that multi-session tSCS in rats with a contusion SCI can be successfully completed across a 6-week experimental timeline to decrease hyperreflexia and increase H-reflex rate modulation. More importantly, we further show that tSCS prevents the SCI-induced disruption in chloride homeostasis in lumbar motoneurons through increasing KCC2 membrane expression, suggesting a way by which tSCS decreases hyperreflexia after SCI (Boulenguez et al., 2010; Côté et al., 2014) and identifying a mechanism through which tSCS contributes to decrease spasticity. Additionally, tSCS increases motor output in ankle muscles after incomplete SCI, as previously reported after complete spinal cord transection (Malloy et al., 2022). This work displays that multi-session tSCS can target and diminish spasticity in a clinically relevant animal model and highlights a return of chloride homeostasis as one contributing factor to the ability of this intervention to prevent the development of spasticity after SCI.

## RESULTS

In a clinically relevant rat model of thoracic contusion SCI, we investigated the effects of tSCS, delivered over 6 weeks (18 sessions), on the development of spasticity and the associated imbalance in chloride homeostasis triggered by SCI. Rats received tSCS 3 times a week for 6 weeks starting 5 days post-SCI. All SCI rats, with (n = 11) and without stimulation (n = 11), received behavioral assessments for functional capabilities (locomotion) and sensory feedback (tactile and thermal) over time. Terminal and post-mortem experiments were performed 6 weeks post-SCI, including age-matched, neurologically intact controls (n = 9), to test spinal reflex excitability and inhibition, motor output in hindlimb muscles, and KCC2 membrane expression in lumbar motoneurons.

### Motor thresholds in response to tSCS over time

We sought to estimate whether motor thresholds in response to tSCS in the awake animal were stable over the18 stimulation sessions. Motor threshold, determined as the lowest intensity to elicit an ankle twitch and measured visually and with light manual touch in the awake animal at rest following one pulse of tSCS, did not fluctuate extensively across sessions and was stable over the first 11 sessions indicating overall consistency in electrode placement across stimulation sessions (**Figure 1**). However, there was a significant decrease (Q = 66.27, p < 0.0001) in motor threshold compared to the first stimulation session at session 12 (p = 0.0028; 79.08 ± 3.87%), session 15 (p = 0.0011; 79.01 ± 3.07%), session 16 (p = 0.0022; 81.88 ± 4.20%), session 17 (p = 0.0033; 80.09 ± 3.84%), and session 18 (p = 0.0185; 82.75 ± 5.40%). This corresponded to the last tSCS session of week four and week five as well as all sessions of week six. Similar changes were also apparent in the absolute stimulus intensity values (in mAmp) as motor threshold (in xMT) significantly decreased in the latter weeks of stimulation starting at session 12 (F9,153 = 24.92, p < 0.0001; range: 2.70 mA; one-way RM ANOVA; *not shown*). This suggests a delayed but potential increase in motor output in response to multi-session tSCS.

**Figure 1.**
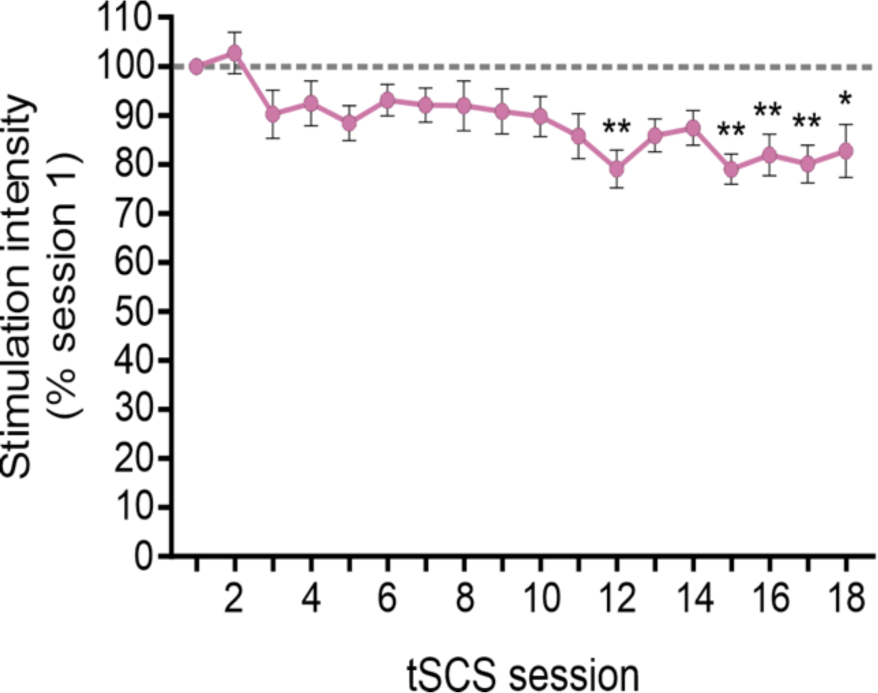
Stimulation intensity required to evoke a motor response in ankle muscles with tSCS in the awake animal. Motor thresholds for SCI + tSCS animals remained relatively consistent across the 18 stimulation sessions but showed a significant decrease in stimulation intensity in the last 5 out of 7 sessions. * p < 0.05; ** p < 0.01; Freidman one-way RM ANOVA. Average ± SEM.

### Multi-session tSCS decreases hyperreflexia after SCI

The H-reflex has long been used as an electrophysiological equivalent of the monosynaptic stretch reflex pathway (Palmieri et al., 2004), and the Hmax/Mmax ratio as well as the low frequency-dependent depression (FDD) of the H-reflex are widely accepted as correlates of hyperreflexia and spasticity, especially in rodents (Thompson et al., 1992; 1998; 2001; Boulenguez et al., 2010). We first evaluated the latency of the motor response in the plantar muscle. Neither SCI nor tSCS affected the onset latency of the M-wave or H-reflex (*not shown*; one-way ANOVA) with similar values in Intact (2.09 ± 0.03 ms, M-wave; 8.54 ± 0.17 ms, H-reflex), SCI (1.94 ± 0.08 ms, M-wave; 8.05 ± 2.08 ms, H-reflex), and SCI + tSCS animals (1.95 ± 0.03 ms, M-wave; 8.14 ± 0.13 ms, H-reflex). To further determine the effect of tSCS on reflex hyperexcitability, we evaluated the rate at which the plantar motoneuronal pool is recruited through tSCS stimulation. M-wave and H-reflex recruitment curves were plotted with peak-to-peak amplitude as a function of increasing stimulus intensities, and a sigmoid function was fitted to the recruitment curve to estimate recruitment properties (**Table 1 & Figure 2**). There were no significant differences between groups for any of the M-wave parameters such as goodness of fit (R^2^), maximal amplitude (Max), slope of the sigmoid function (m), stimulus intensity at 50% of maximal amplitude (S50), stimulus intensity to reach threshold (Threshold; mA), and intensity required to reach maximal amplitude (x MT) (**Table 1, top**).

**Figure 2.**
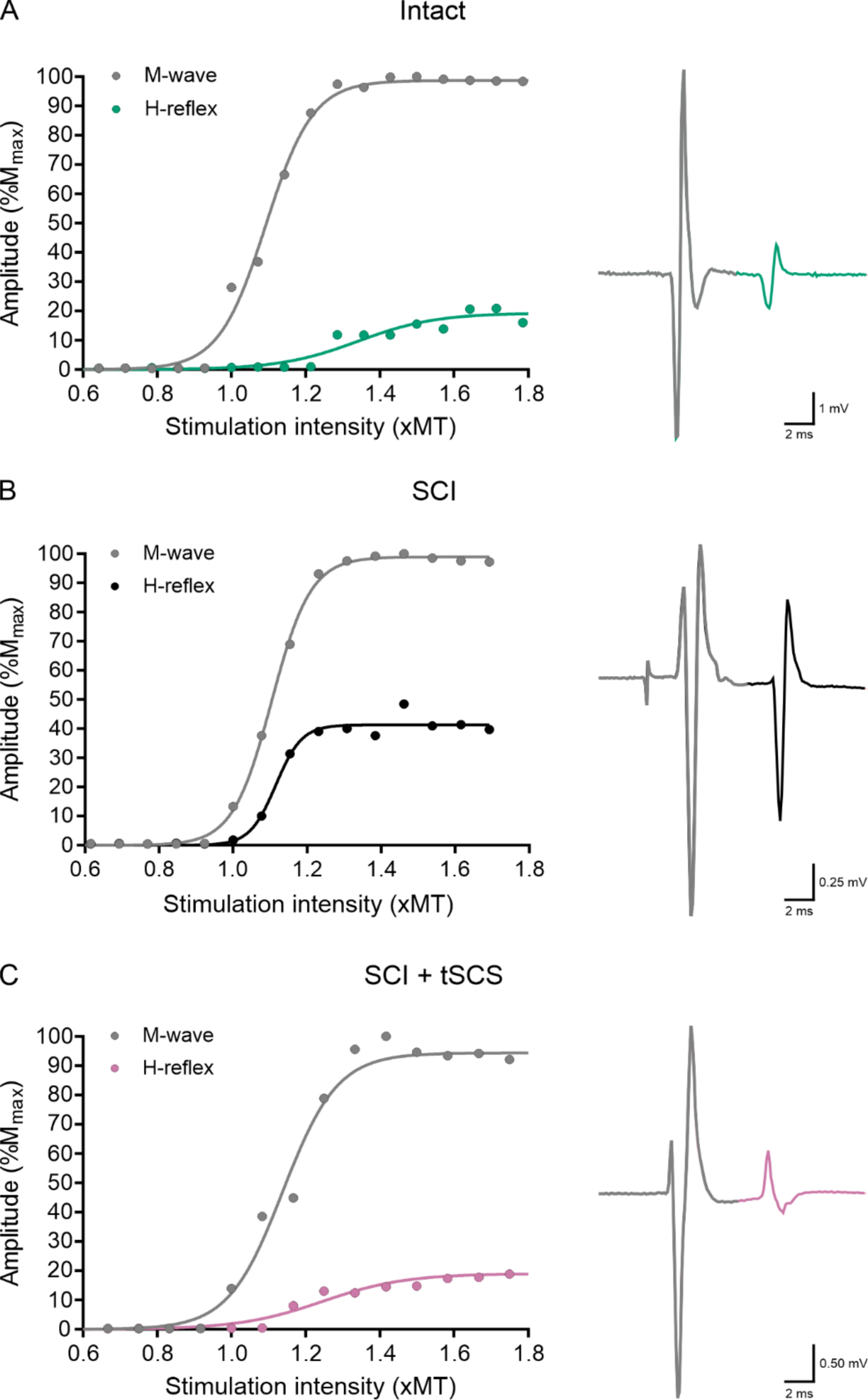
Representative examples of M-wave and H-reflex recruitment curves. Individual recruitment curves (left) and representative traces (right, 1.4 xMT) for (**A**) Intact (**B**) SCI, and (**C**) SCI + tSCS animals show no difference in M-wave recruitment properties but identifiable differences in maximal H-reflex amplitude, H-reflex threshold intensity, and slope of the H-reflex sigmoid function.

**Table 1.** Parameters of sigmoid function for the M-wave and H-reflex recruitment curves.

| Waveform | Group | $R^2$ | Max amplitude (mV) | $m$ | S50 (× MT) | Threshold (mA) | Intensity to reach Max (× MT) |
| --- | --- | --- | --- | --- | --- | --- | --- |
| M-wave | Intact | $0.97 \pm 0.022$ | $2.88 \pm 0.73$ | $12.91 \pm 4.40$ | $1.39 \pm 0.22$ | $0.029 \pm 0.009$ | $1.78 \pm 0.44$ |
| | SCI | $0.99 \pm 0.002$ | $4.21 \pm 1.06$ | $14.63 \pm 2.08$ | $1.81 \pm 0.05$ | $0.037 \pm 0.023$ | $1.36 \pm 0.08$ |
| | SCI + tSCS | $0.98 \pm 0.003$ | $3.62 \pm 0.50$ | $17.11 \pm 1.73$ | $1.18 \pm 0.05$ | $0.056 \pm 0.051$ | $1.31 \pm 0.06$ |
| H-reflex | Intact | $0.91 \pm 0.021$ | **$0.47 \pm 0.14^{**}$** | $16.75 \pm 3.33$ | $1.67 \pm 0.35$ | $0.039 \pm 0.001$ | $1.77 \pm 0.33$ |
| | SCI | $0.97 \pm 0.018$ | **$2.69 \pm 0.92^{**/###}$** | **$26.81 \pm 5.24^{\#}$** | $1.27 \pm 0.06$ | $0.037 \pm 0.011$ | $1.37 \pm 0.06$ |
| | SCI + tSCS | $0.94 \pm 0.018$ | **$0.72 \pm 0.22^{\#}$** | **$12.05 \pm 1.52^{\#}$** | $1.26 \pm 0.08$ | $0.051 \pm 0.020$ | $1.37 \pm 0.10$ |

Average ± SEM predicted values from the sigmoid fit with stimulation intensities plotted against response amplitudes. $R^2$ is the goodness of fit; Max amplitude (mV) is the maximal amplitude of the response; $m$ is the slope parameter of the sigmoid function; S50 ( × MT) is the stimulus intensity required to elicit a response equivalent to 50% of the maximal amplitude; Threshold (mA) and Intensity to reach Max amplitude ( × MT) are stimulation intensities corresponding to threshold and maximal amplitudes, respectively. Bolded values denote significant differences between groups. \*, Intact vs SCI; #, SCI vs SCI + tSCS; # $p < 0.05$ ; \*\*/### $p < 0.01$ ; Kruskal-Wallis one-way ANOVA.

For the H-reflex, no significant differences were found for R^2^, S50, Threshold, or Intensity to reach Max amplitude between groups (**Table 1, bottom**). However, maximal H-reflex amplitude (Hmax) was significantly different between groups (**Table 1**; H (2) = 9.802, p = 0.007) with SCI animals showing increased Hmax amplitude compared to Intact (p = 0.014) and SCI + tSCS groups (p = 0.021). This is also apparent in both individual (**Figure 2**) and group averaged (**Figure 3**) recruitment curves. Stimulation intensity (mA) required to evoke a response (threshold) of the H-reflex was not significantly different between groups (**Table 1**); however, it was significantly greater for Intact versus SCI and SCI + tSCS animals when normalized to motor threshold (**Figure 2**; H (2) = 10.537, p = 0.005, Kruskal-Wallis one-way ANOVA). Further, the slope of the sigmoid function (m) was significantly decreased for SCI + tSCS compared to SCI animals (**Table 1**; H (2) = 6.43, p < 0.05). But the SCI group was not statistically different from the Intact group (**Table 1**). Individual animal Hmax/Mmax ratios were also significantly different across groups (**Figure 2**, H (2) = 10.157, p = 0.006, Kruskal-Wallis one-way ANOVA) showing a substantial increase in SCI compared to Intact animals (p = 0.043) and SCI + tSCS (p = 0.007).

**Figure 3.**
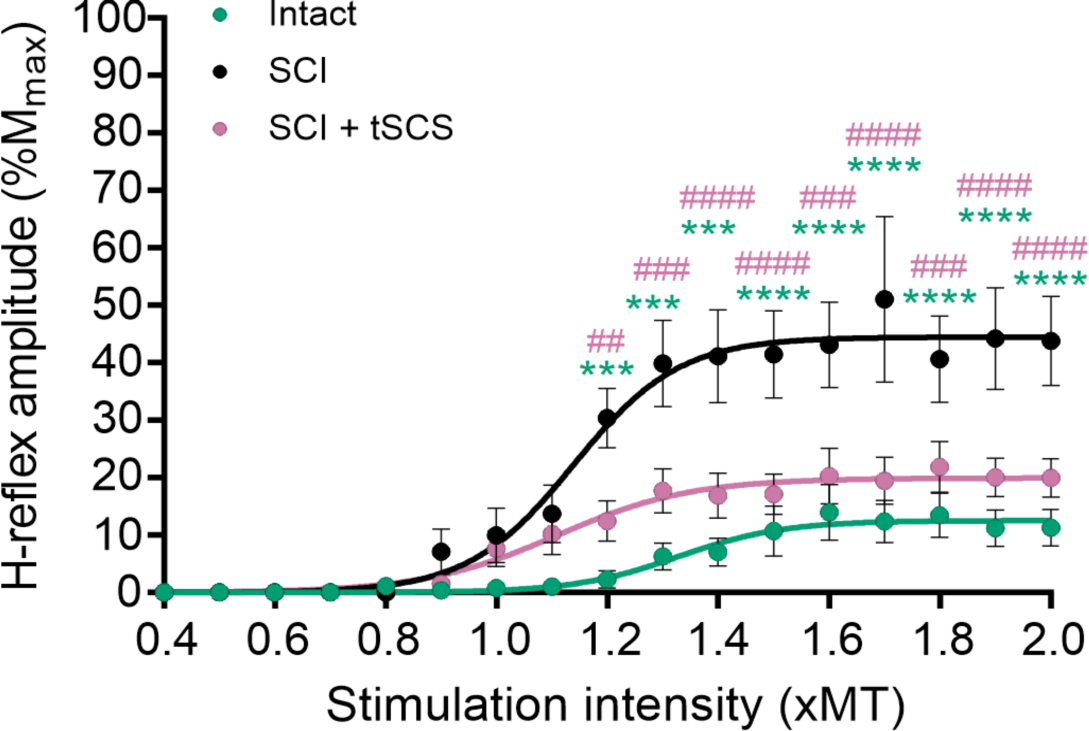
Multi-session tSCS after SCI decreases hyperreflexia. Group average recruitment curves of the H-reflex show SCI animals with an H-reflex amplitude significantly greater than the Intact and SCI + tSCS groups while the latter have no differences between each other. *, Intact vs SCI; #, SCI vs SCI + tSCS. **/## p < 0.01; ***/### p < 0.001; ****/#### p < 0.0001; two-way RM ANOVA. Average ± SEM.

Normalized values from individual recruitment curves were used to establish a group sigmoid function. A two-way RM ANOVA revealed significant differences in H-reflex amplitudes between groups (F2,21 = 9.876, p = 0.0009) and at stimulation intensities (F17,357 = 29.65, p < 0.0001). There was also a significant interaction between groups and stimulation intensity for H-reflex amplitudes (**Figure 3**; F34,357 = 5.529, p < 0.0001). Overall, Holm-Sidak post hoc multiple comparisons test indicated that SCI animals showed significantly increased H-reflex amplitudes at intensities ranging from 1.2 to 2.0 xMT compared to the Intact and SCI + tSCS groups (p < 0.01), which were not different from each other. Group Hmax/Mmax ratios further showed significant increases in SCI compared to Intact animals which was improved by tSCS as these animals were not significantly different from Intact (**Figure 3**). Thus, these results suggest that tSCS can decrease overall hyperreflexia of the plantar monosynaptic reflex pathway observed in SCI animals.

### Multi-session tSCS improves H-reflex reflex rate modulation after SCI

We further evaluated the effect of tSCS on H-reflex modulation by analyzing the frequency dependent depression (FDD). H-reflex amplitude was compared between groups across increasing stimulus frequencies. A two-way RM ANOVA revealed significant differences in H-reflex amplitudes between stimulation frequencies (F2,44 = 346, p < 0.0001) but not between groups. However, there was a significant interaction between groups and frequency of stimulation for H-reflex amplitude (**Figure 4**; F4,44 = 2.911, p = 0.032). Holm-Sidak post hoc multiple comparisons test revealed SCI animals with significantly greater H-reflex amplitude at 5 Hz stimulation (31.83 ± 11.13%) compared to Intact (p = 0.0058; 0.69 ± 0.52%) and SCI + tSCS (p = 0.0406; 13.44 ± 3.36%) animals, which indicates an impairment of reflex modulation. The SCI group also showed greater H-reflex amplitude at 10 Hz stimulation (24.37 ± 10.83%) compared to the Intact (0.55 ± 0.55%) and SCI + tSCS (12.31 ± 3.22%) groups, but this was only statistically significant versus Intact animals (p = 0.0477), likely due to large variability within the SCI group. This indicates that multi-session tSCS after SCI can, at least partially, restore H-reflex modulation in a frequency dependent manner similarly to Intact animals.

**Figure 4.**
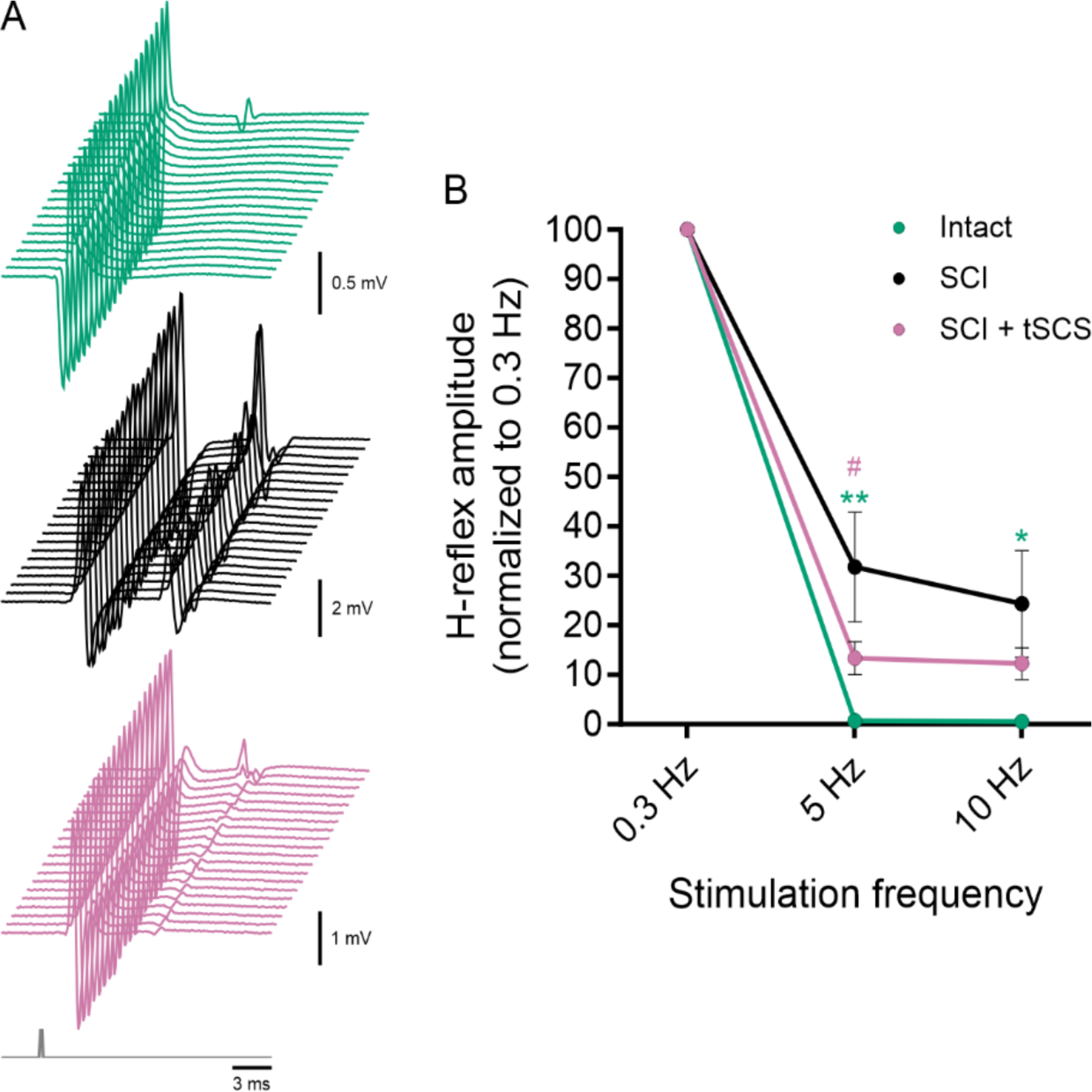
Frequency dependent depression of the H-reflex. Rate modulation of the H-reflex was assessed by measuring the H-reflex amplitude in response to consecutive tibial nerve stimulations at various increasing intensities. (**A**) Representative traces of M-wave and H-reflex responses to 5 Hz stimulation between Intact (green), SCI (black), and SCI + tSCS (pink) groups. (**B**) SCI animals showed significantly less depression of H-reflex amplitude at 5 and 10 Hz compared to Intact animals. The SCI + tSCS group showed a significant increase in rate modulation at 5 Hz but not 10 Hz compared to the SCI group. There were no differences between Intact and SCI + tSCS animals across stimulation frequencies. *, Intact vs SCI; #, SCI vs SCI + tSCS. */# p < 0.05; ** p < 0.01; two-way RM ANOVA. Average ± SEM.

### Multi-session tSCS prevents KCC2 membrane downregulation in lumbar motoneurons after SCI

Previous work within our laboratory indicates that the repetitive activation of the neuromuscular system with exercise after SCI prevents the deterioration of the FDD of the H-reflex by counteracting the SCI-induced decrease in KCC2 expression on the motoneuron membrane (Beverungen et al., 2020; Côté et al., 2014). We evaluated whether tSCS similarly prevents the downregulation of KCC2 membrane expression after SCI. **Figure 5A** shows representative images of KCC2 immunolabelling on ChAT^+^ lumbar motoneurons between Intact, SCI, and SCI + tSCS animals. The membrane to cytosol ratio of KCC2 fluorescence expression was significantly lower in the SCI group (1.13 ± 0.03) compared to Intact (2.44 ± 0.12) and SCI + tSCS (2.08 ± 0.11) animals (**Figure 5B**; H (3) = 67.9, p < 0.0001), which were not different from each other. This reduction in KCC2 membrane expression following SCI was prevented by tSCS with values similar to Intact, resembling what has been reported with activity-based therapies (Beverungen et al., 2020; Bilchak et al., 2021; Chopek et al., 2015; Côté et al., 2014; Tashiro et al., 2015).

**Figure 5.**
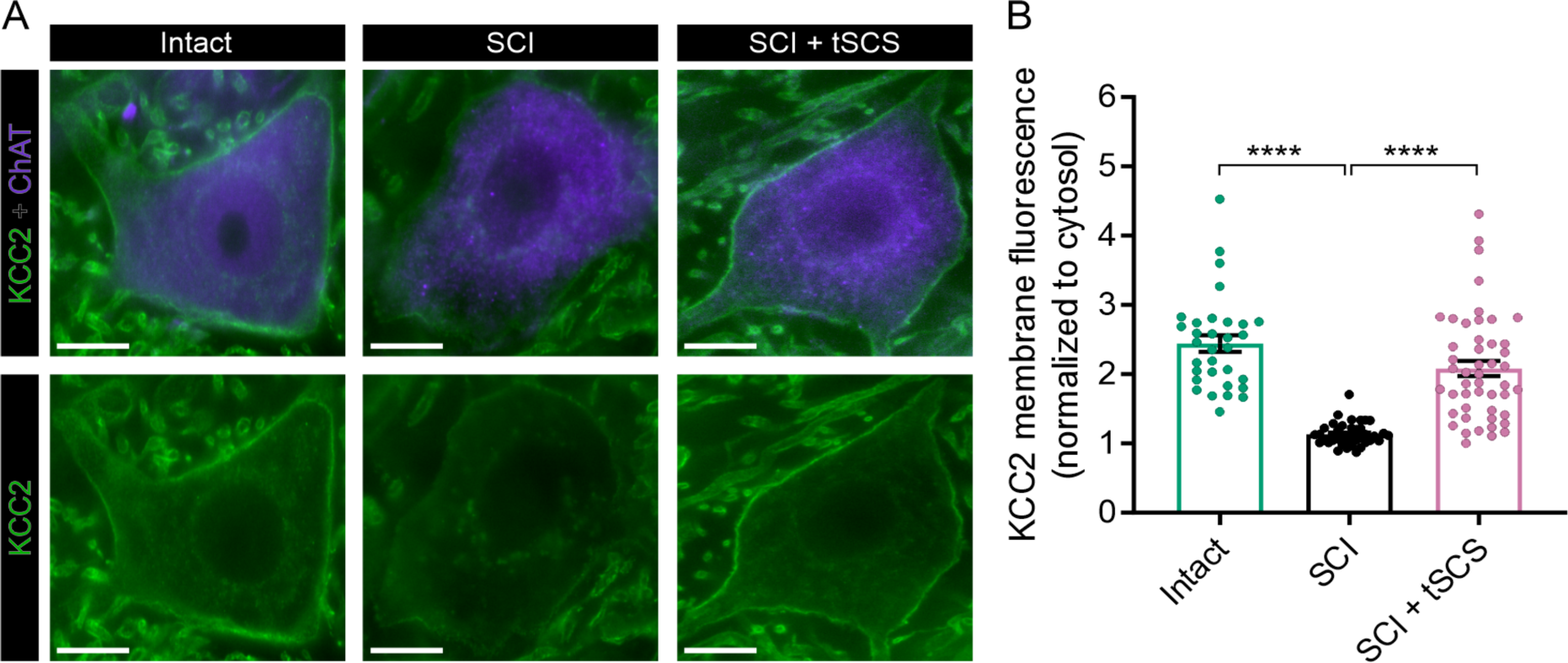
KCC2 membrane expression in lumbar motoneurons. (**A**) Representative images of KCC2 immunolabelling in ChAT^+^ motoneurons in the lumbar ventral horn between Intact, SCI, and SCI + tSCS animals. (**B**) Quantification of motoneuronal KCC2 membrane fluorescence normalized to cytosolic expression. The SCI group showed a significantly decreased expression of KCC2 on the motoneuronal membrane compared to Intact and SCI + tSCS animals which were not different from each other. KCC2, potassium-chloride cotransporter isoform 2; ChAT, choline acetyl transferase. Scale bar: 10 µm. **** p < 0.0001; Kruskal-Wallis one-way ANOVA. Average ± SEM.

### Multi-session tSCS increases motor output in ankle extensor muscles after SCI

Additionally, we evaluated spinal excitability and maximal output of hindlimb motor pools in response to tSCS by analyzing recruitment properties. Intact, SCI, and SCI + tSCS animals showed no difference in response onset latency for MG (2.74 ± 0.10 ms, Intact; 2.50 ± 0.06 ms, SCI; 2.74 ± 0.11 ms, SCI + tSCS) or TA (2.70 ± 0.09 ms, Intact; 2.60 ± 0.11 ms, SCI; 2.89 ± 0.13 ms, SCI + tSCS) muscles (*not shown*; one-way ANOVA). Also, there were no significant differences in transspinal evoked potential (TEP) recruitment curve parameters such as goodness of fit (R^2^), maximal amplitude (Max), slope of the sigmoid function (m), stimulus intensity at 50% of maximal amplitude (S50), slope of the sigmoid function confined to occur at S50 (Slope), and stimulus intensity (xMT) for maximal amplitude (Max) between groups for MG or TA muscles (**Table 2**). SCI animals required a significantly higher stimulus intensity (Threshold) to evoke a response in the TA (F2,23 = 6.379, p = 0.0063) but not MG compared to Intact animals (**Table 2, bottom**). SCI + tSCS animals did not show any significant differences between groups for Threshold in the MG or TA.

**Table 2.** Parameters of sigmoid function for the transspinal evoked potential (TEP) recruitment curves in ankle muscles.

| Muscle | Group | R<sup>2</sup> | Max amplitude (mV) | m | S50 (x MT) | Slope | Threshold (mA) | Intensity to reach Max (x MT) |
| --- | --- | --- | --- | --- | --- | --- | --- | --- |
| MG | Intact | 0.97 ± 0.004 | 13.60 ± 1.99 | 4.32 ± 0.99 | 1.63 ± 0.11 | 0.63 ± 0.11 | 3.16 ± 0.27 | 2.26 ± 0.22 |
|  | SCI | 0.98 ± 0.005 | 23.32 ± 5.12 | 5.61 ± 0.82 | 1.42 ± 0.06 | 0.42 ± 0.06 | 4.33 ± 0.32 | 1.84 ± 0.12 |
|  | SCI + tSCS | 0.99 ± 0.003 | 12.80 ± 2.89 | 9.18 ± 2.08 | 1.31 ± 0.09 | 0.31 ± 0.09 | 3.61 ± 0.45 | 1.62 ± 0.18 |
| TA | Intact | 0.98 ± 0.004 | 12.72 ± 2.86 | 3.82 ± 0.59 | 1.65 ± 0.11 | 0.65 ± 0.11 | **2.81 ± 0.41\*** | 2.29 ± 0.22 |
|  | SCI | 0.99 ± 0.002 | 12.12 ± 2.49 | 6.21 ± 0.89 | 1.38 ± 0.06 | 0.38 ± 0.06 | **5.08 ± 0.28\*** | 1.75 ± 0.12 |
|  | SCI + tSCS | 0.99 ± 0.004 | 12.72 ± 2.88 | 5.59 ± 0.83 | 1.46 ± 0.09 | 0.46 ± 0.09 | 3.97 ± 0.57 | 1.91 ± 0.18 |

Average ± SEM predicted values from the sigmoid fit with stimulation intensities plotted against response amplitudes. R<sup>2</sup> is the goodness of fit; Max amplitude (mV) is the maximal amplitude of the response; m is the slope parameter of the sigmoid function; S50 (xMT) is the stimulus intensity required to elicit a response equivalent to 50% of the maximal amplitude; Slope is the slope of the sigmoid relationship confined to occur at S50; Threshold (mA) and Intensity to reach Max amplitude (xMT) are stimulation intensities corresponding to threshold and maximal amplitudes, respectively. MG, medial gastrocnemius; TA, tibialis anterior. Bolded values denote significant differences between groups. \*, Intact vs SCI; \* p < 0.05; one-way ANOVA.

Two-way RM ANOVAs revealed significant differences in TEP amplitudes in the MG and TA muscles between groups (F2,20 = 6.164, p = 0.0082, MG; F2,22 = 3.442, p = 0.05, TA) and at stimulation intensities (F16,320 = 244.9, p < 0.0001, MG; F16,352 = 295.6, p < 0.0001, TA). While tSCS did not affect the intensity required to evoke a response (Threshold) in the MG, SCI + tSCS animals did show a significant interaction between groups (vs. Intact, SCI) and stimulation intensity through increased relative amplitude for TEPs in the MG (F32,320 = 4.823, p < 0.0001) at intensities ranging from 1.2 to 2.0 xMT compared to Intact and SCI groups (**Figure 6A**; p < 0.05, Holm-Sidak post hoc). Similarly, SCI animals also showed a significant interaction between groups (vs. Intact) and stimulation intensity with an increase in MG amplitude compared to Intact animals at intensities ranging from 1.3 to 1.9 xMT (F32,320 = 4.823, p < 0.0001), but multiple comparisons with Holm-Sidak post hoc test indicated that this remained significantly smaller than the SCI + tSCS group (**Figure 6A**; p < 0.05). Like the MG, there was a significant interaction between groups (vs. Intact) and stimulation intensity as TEP amplitudes were also increased in TA in both SCI groups as compared to Intact animals from 1.2 to 2.0 xMT, but there was no additional effect of tSCS identified through Holm-Sidak post hoc test compared to the SCI group **(Figure 6B**; F32,352 = 2.677, p < 0.0001). This suggests that tSCS increases the recruitment gain as well as the overall motor output after SCI with a potential preferential effect on ankle extensor versus flexor muscles. Furthermore, tSCS did not negatively impact locomotor capabilities as BBB scores for SCI (8.50 ± 0.45; 6 weeks post-SCI) and SCI + tSCS (9.92 ± 0.39; 6 weeks post-SCI) animals were not statistically different across 2-, 4-, or 6-week post-SCI timepoints (*not shown*; two-way RM ANOVA). Thus, indicating that this protocol of tSCS will not impair locomotor capacity despite not being optimized to specifically target this functional task.

**Figure 6.**
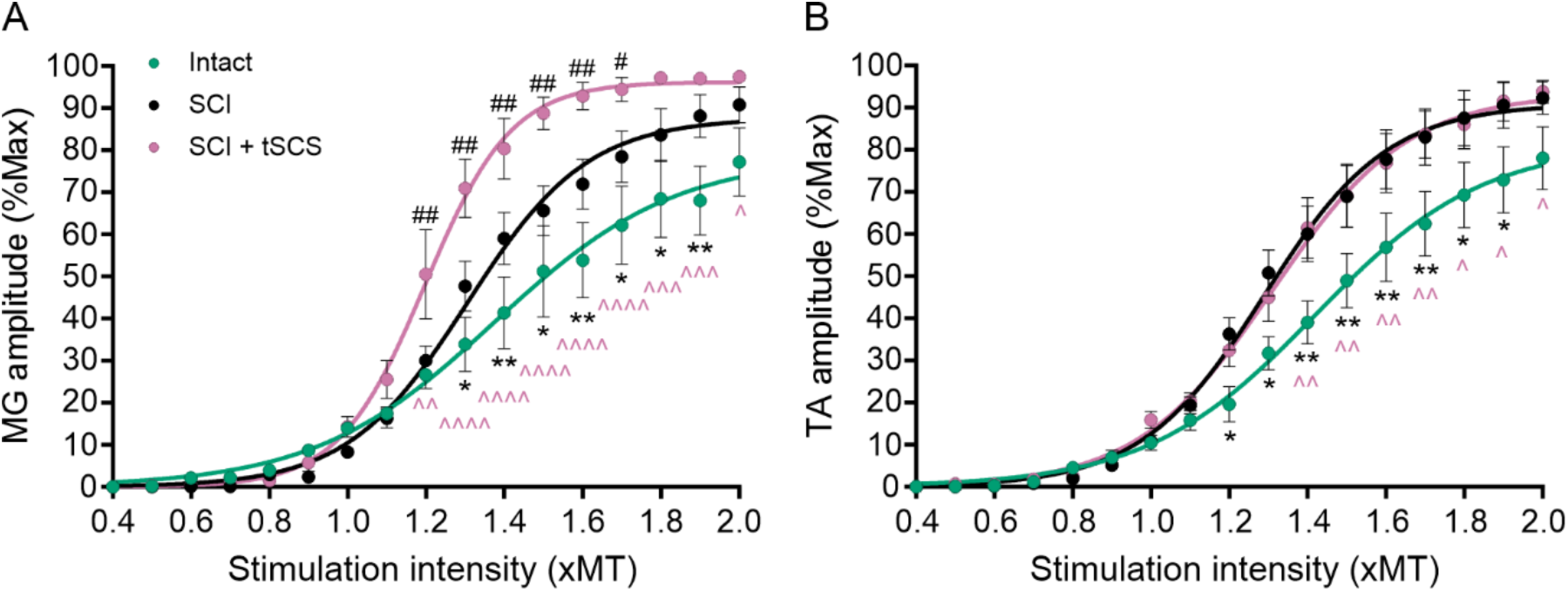
Multi-session tSCS after SCI increases motor output in ankle extensor muscles. The recruitment curves of TEPs in MG (**A**) and TA (**B**) muscles show significantly increased amplitudes at various intensities (xMT) for SCI compared to Intact animals. This response is identical for the SCI + tSCS compared to SCI group in the TA but is augmented in the MG. MG, medial gastrocnemius; TA, tibialis anterior. *, Intact vs SCI; ^, Intact vs SCI + tSCS; #, SCI vs SCI + tSCS. */^/# p < 0.05; **/^^/## p < 0.01; ^^^ p < 0.001; ^^^^ p < 0.0001; two-way RM ANOVA. Average ± SEM.

### Multi-session tSCS does not induce maladaptive hypersensitivity after SCI

We then tested the effect of tSCS on paw withdrawal thresholds to tactile and thermal stimuli. Tactile stimulus with DVFH tests (**Figure 7A**) revealed no significant differences for withdrawal thresholds between groups. However, within groups, there were significant differences between timepoints as both groups showed decreased withdrawal thresholds compared to baseline, suggesting that SCI negatively impacts withdrawal responses to tactile sensation (F4,80 = 6.363, p = 0.0002). There was no interaction with groups over time as there were not significant differences between groups for paw withdrawal threshold at 1 week post-SCI (104.91 ± 20.35%, SCI; 72.27 ± 10.31%, SCI + tSCS), 2 weeks post-SCI (74.64 ± 12.31%, SCI; 64.73 ± 6.33%, SCI + tSCS), 4 weeks post-SCI (72.46 ± 8.61%, SCI; 59.91 ± 4.09%, SCI + tSCS), or 6 weeks post-SCI (86.36 ± 5.61%, SCI; 67.36 ± 8.78%, SCI + tSCS). Hargreaves was analyzed starting 5 weeks post-SCI when most animals were considered weight-bearing on both hindlimbs. There were insufficient sample sizes within groups to statistically analyze timepoints earlier than this, as Hargreaves requires bilateral weight-bearing ability to perform the assessment. There were no significant differences between groups or timepoints and no interaction between groups at 5- and 6-weeks post-SCI for paw withdrawal in response to thermal stimulus **(Figure 7B**; 91.63 ± 5.64%, SCI; 102.4 ± 9.63%, SCI + tSCS). Collectively, this indicates that multi-session tSCS after SCI does not induce maladaptive hypersensitivity related to developing mechanical allodynia or thermal hyperalgesia.

**Figure 7.**
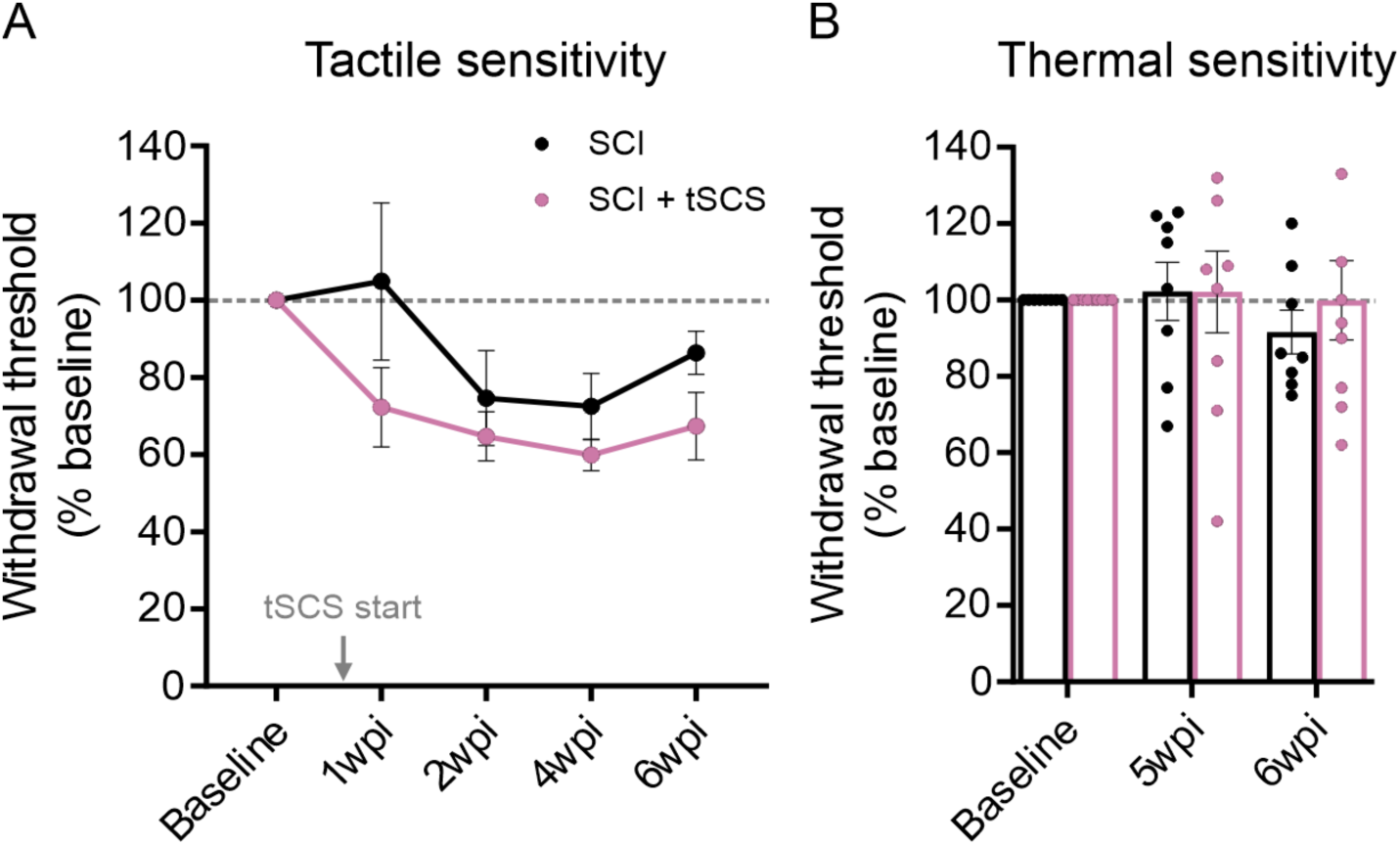
Tactile and thermal sensory testing over time. (**A**) Dorsal von Frey hair was used to measure withdrawal threshold latency to tactile sensation across timepoints. Latency to paw withdrawal was not significantly different between SCI and SCI + tSCS animals. (**B**) Hargreaves was used to measure withdrawal threshold latency to thermal sensations across timepoints. There were no differences between groups at 5- and 6-weeks post-SCI (n=8/group). p > 0.05; two-way Mixed ANOVA. Average ± SEM.

## DISCUSSION

Spasticity is difficult to manage, and while current pharmacological approaches such as anti-spastic medication is the most common therapeutic approach, its success rate is low (Taricco et al., 2006) and only 38% of individuals with SCI report an improvement in spastic symptoms (Field-Fote et al., 2022). Indeed, antispasmodics induce a pronounced depression in overall neural excitability, thereby leading to reduced muscle activity and weakness, which impedes rehabilitation efforts, the potential of restoring voluntary movements, and the performance of activities of daily living (Angeli et al., 2012; Dario and Tomei, 2004; Dietz et al., 2023; Elovic, 2001; Kirshblum, 1999). It is often further accompanied by unpleasant side effects such as impaired memory, dizziness, and sedation (Dario and Tomei, 2004; Dietz et al., 2023; Elovic, 2001; Gracies et al., 1997; Kirshblum, 1999). These undesirable effects critically affect quality of life for people with SCI, so identifying alternatives to pharmacological therapeutics for improving spasticity and elucidating the mechanisms at play remains critical. Our results suggest that multi-session tSCS is a viable alternative approach that not only successfully improves hyperreflexia but does not indiscriminately dampen motor output.

The interest toward tSCS in the context of SCI is recent and has focused on functional improvements after decades of research mainly dedicated to treating pain (Reviewed in Caylor et al., 2019; Lam et al., 2023; Tapias Perez, 2022). Recent reports in humans suggest that noninvasive tSCS after SCI not only has the potential to improve motor recovery, including walking (Megia Garcia et al., 2020; Seanez and Capogrosso, 2021) and hand function (Inanici et al., 2018; 2021), but also to potentially act as an anti-spastic therapeutic strategy. While the immediate effect of a single session of tSCS (15-30 min.) on alleviating spasticity in an acute setting is encouraging and plentiful (Estes et al., 2018; 2017; Hofstoetter et al., 2014; 2020; Sandler et al., 2021), the long-term effects of multi-session tSCS are currently being studied (Knikou and Murray, 2019; Murray and Knikou, 2019). We show here, for the first time, that the positive effects of multi-session tSCS on spasticity are indicative of structural and physiological plastic changes within the spinal cord, and we have identified that one of the underlying mechanisms include prevention of a disruption in chloride homeostasis induced by SCI.

### Successfully targeting SCI-induced hyperreflexia with multi-session tSCS

We used a newly developed, clinically relevant animal model with an incomplete SCI receiving tSCS in an awake setting (Malloy et al., 2022), and we probed spinal reflex excitability, neuroanatomical plasticity, and functional recovery following 6 weeks of stimulation (18 sessions). Spinal reflex excitability was assessed via the H-reflex, which has long been used as an electrophysiological equivalent of the monosynaptic stretch reflex pathway (Palmieri et al., 2004). The amplitude of the H-reflex is an estimation of the number of motor units activated, and any modulation indicates a change in the excitability in this pathway. In addition, the Hmax/Mmax ratio as well as the low frequency-dependent depression (FDD) of the H-reflex are well-established correlates of hyperreflexia and spasticity after SCI both in rodents (Boulenguez et al., 2010; Skinner et al., 1996; Thompson et al., 1992) and humans (Calancie et al., 1993; Ishikawa et al., 1966).

We found that tSCS significantly decreased both the Hmax and the Hmax/Mmax ratio as well as reduced the slope of the recruitment curve of the H-reflex toward Intact values. We further observed a significant improvement in the FDD of the H-reflex following 5 Hz stimulation, indicating an increase in rate modulation of the plantar monosynaptic reflex pathway. While tSCS did not significantly improve FDD at 10 Hz stimulation compared to control SCI animals, the variability seen in the SCI group is common as rats often display a wide range of hyperreflexia after contusion injury (Petrosyan et al., 2020, see also **Figure 4**). The extent of variability in SCI animals may be a result of the extensive ability of rats to recover locomotion and functional capabilities after contusion SCI despite minimal white matter sparing (Magnuson et al., 2005; Scheff et al., 2003). Additionally, spasticity in humans is prevalent in around 75% of individuals with SCI (Holtz et al., 2017; Maynard et al., 1990), so it is equally likely that only a similar proportion of our animals truly developed spasticity after injury. Since widely accepted behavioral or noninvasive measurements for the development of spasticity in rodents are lacking, this remains a limitation of our investigations. However, the tight variability in the SCI + tSCS group suggests that this is a true group effect and that tSCS can promote functional recovery to an equal extent across animals.

Together, our results suggest that tSCS decreases hyperreflexia after SCI. This is further supported by recent literature with human-based tSCS where H-reflex amplitude was significantly decreased following stimulation, as per recruitment curves, while M-wave properties remained unchanged, and homosynaptic depression of H-reflex amplitude was apparent (Knikou and Murray, 2019). Both of which are akin to our results investigating recruitment properties and low frequency-dependent depression of the H-reflex in rat. The decrease in H-reflex excitability observed following SCI and restoration with tSCS could be attributed to both synaptic and non-synaptic plasticity. Low frequency-dependent depression is a gradual decrease in H-reflex amplitude that occurs in response to a train of stimuli at frequencies between 1 and 10 Hz (Lloyd and Wilson, 1957). Mechanisms responsible for impaired low frequency-dependent depression after SCI include a disruption in presynaptic inhibition of primary afferent terminals and/or impaired homosynaptic depression (Calancie et al., 1993; Delwaide, 1973; Eccles and Rall, 1951; Hultborn et al., 1996; Kohn et al., 1997; Lloyd and Wilson, 1957; Thompson et al., 1992). Our results further suggest that multi-session tSCS potentially contributes to prevent the SCI-induced maladaptive plasticity in these pathways.

### Activity-based therapies: commonalities between tSCS and exercise suggest similarities in underlying mechanisms

Another alternative to pharmacological therapeutics to decrease spasticity, and perhaps the most successful according to individuals with SCI, is exercise-based rehabilitation (Field-Fote et al., 2022). This includes locomotor training, which is not only associated with improved spastic symptoms following SCI (Adams and Hicks, 2011; Manella and Field-Fote, 2013), but even more so than anti-spastic medication (Mirbagheri, 2015). Importantly, reduced spasticity in response to exercise is thought to be driven by the activation of sensory afferents (Fong et al., 2009; Rossignol and Frigon, 2011; Smith and Knikou, 2016). tSCS also engages sensory afferents of large and medium diameter in the lumbar dorsal roots (Danner et al., 2011; Hofstoetter et al., 2018; Minassian et al., 2011), and growing evidence suggests it could contribute to reduce spasticity after SCI (Estes et al., 2017; Hofstoetter et al., 2014; 2020). This provides a framework suggesting that spinal cord stimulation and exercise-based rehabilitation may share common elements and mechanisms. The decrease we observe in Hmax amplitude, reduced Hmax/Mmax ratio, and improved FDD further support the similarity between the effect of tSCS and exercise. Indeed, we and others have shown that passive cycling and step-training decrease the Hmax/Mmax ratio as well as the slope of the H-reflex recruitment curve and improve the FDD of the H-reflex and post-synaptic inhibition in trained versus unexercised animals after SCI (Caron et al., 2020; Côté et al., 2011; Reese et al., 2006; Tashiro et al., 2015). While a more detailed investigation is needed, it has been hypothesized that afferent inputs of large and medium diameter directly activate spinal interneurons (Côté et al., 2018; Jankowska, 1992), and this effect likely contributes to the improvements in spastic symptoms after SCI that are associated with activity-based therapies, including various forms of afferent stimulation.

#### Multi-session tSCS prevents the SCI-induced decrease in motoneuronal KCC2

Previous work within our laboratory has shown that improving FDD of the H-reflex with activity-based therapies is associated with a brain-derived neurotrophic factor (BDNF)-induced increase in KCC2 expression on the motoneuron membrane, thereby contributing to decreased hyperreflexia after SCI (Beverungen et al., 2020; Bilchak et al., 2021; Chopek et al., 2015; Côté et al., 2014; Tashiro et al., 2015). KCC2, a chloride extrusion transporter, is required to keep the intracellular concentration of chloride low, thereby maintaining the equilibrium potential for chloride low (farther away from the action potential threshold) and allowing for hyperpolarizing inhibitory postsynaptic potentials (Delpire and Mount, 2002). After SCI, the internalization of KCC2 from the motoneuronal membrane increases the intracellular concentration of chloride, thereby shifting the equilibrium potential for chloride high (toward the action potential threshold) and causing inhibitory postsynaptic potentials to either lose strength of inhibition or switch toward facilitation (Boulenguez et al., 2010). This creates a decrease in postsynaptic inhibition of the motoneuron, which has been strongly implicated in reflex disinhibition and spasticity (Boulenguez et al., 2010; Gackiere and Vinay, 2015; Lu et al., 2008; Rivera et al., 2002). Here, we found that tSCS also significantly increases the motoneuronal membrane expression of KCC2 to near Intact levels, indicating a restoration of motoneuronal chloride homeostasis. Our findings, again, suggest a likely connection between the mode of these two interventions and their mechanisms of plasticity. While the mechanisms by which activity-based therapies modulate chloride cotransporters remain to fully elucidate (Reviewed in Côté, 2020), the importance of an activity-dependent increase in BDNF has been established (Beverungen et al., 2020). Additionally, electrical stimulation has been shown to increase neuronal BDNF expression and upregulate its primary receptor target, tropomyosin receptor kinase B (TrkB), indicating that electrical stimulation can activate the BDNF-TrkB signaling pathway (Dorrian et al., 2023; Ghorbani et al., 2020; McGregor and English, 2018). As both interventions function by repetitively activating sensory afferents, this suggests that tSCS likely acts similarly to activity-based therapies to restore motoneuronal chloride homeostasis and may be BDNF-dependent. Further studies will be necessary to confirm this.

#### Multi-session tSCS preferably increases motor output in extensor muscles

Beyond improved spasticity, our results further suggest an increase in motor output, as illustrated by an increase in TEP amplitude after 6 weeks of tSCS in contused animals, which is similar to what was observed in our complete transected model (Malloy et al., 2022). This was also recently reported in one study with humans receiving a similar protocol of tSCS (Murray and Knikou, 2019). TEPs were shown to be initiated in the posterior roots in humans (Angeli et al., 2014; Calvert et al., 2019; Hofstoetter et al., 2008; Hofstoetter et al., 2014; Minassian et al., 2007a; Minassian et al., 2011; Minassian et al., 2016b; Sayenko et al., 2019; Wagner et al., 2018) and animals (Malloy et al., 2022; Mishra et al., 2017) and are generally believed to result from the activation of proprioceptive afferents within the dorsal roots which leads to a synchronized response of lumbar motoneurons (Capogrosso et al., 2013; Danner et al., 2011, Ladenbauer et al 2010, Rattay et al., 2000,). During movement, tSCS can facilitate motor output to enable standing and stepping as well as voluntary movement of the legs (Angeli et al., 2014; Gerasimenko et al., 2015c; Hofstoetter et al., 2017; Rejc et al., 2017; Sayenko et al., 2019) and arms/hands (Gad et al., 2018; Inanici et al., 2018; 2021). However, these effects of tSCS on motor output are in the context of active stimulation (i.e., stimulation is “on” during movement). This immediate beneficial effect of spinal cord stimulation on motor output is believed to result from the summation of weak, subthreshold, volitional descending signals that increase the excitability of spinal networks and bring motoneurons to threshold. But definitive evidence for the mechanisms by which spinal cord stimulation induces long-term plastic changes in the spinal cord without active stimulation (i.e., when the stimulator is “off”), such that is observed here following weeks of stimulation, remains to be determined. Nevertheless, multi-session tSCS not only decreased H-reflex hyperexcitability, but also amplified the responsiveness of motoneurons as suggested by the leftward shift of the TEP recruitment curves.

Interestingly, increased motor output in the tSCS group favored extensor muscles in our contusion injury model as only significant increases were reported in the MG but not the TA muscle, unlike after a complete spinal transection (Malloy et al., 2022). Since both SCI and SCI + tSCS groups showed significantly increased motor output and gain of recruitment compared to Intact animals, it is likely that part of this shift in excitability is attributed to local plasticity induced by the injury, but tSCS further augmented this shift in ankle extensor muscles. This suggests a role for the spared supraspinal tracts in initiating spinal plasticity in the incomplete injury model. This is further supported by previous human-based investigations with tSCS also showing mixed effects of lower extremity motor output between SCI individuals with American Spinal Injury Association Impairment Score (AIS) A-B versus AIS C-D where AIS C-D individuals also showed a preferential motor increase in ankle extensors but not flexors (Murray and Knikou, 2019). Furthermore, the pro-extensor effect of tSCS is also supported by an increased amplitude of motor responses evoked in the quadriceps muscles but not in the TA immediately after tSCS in healthy people (Megia-Garcia et al., 2020). Potential differences in input from spared descending projections onto extensor versus flexor muscles in contused SCI animals will need to be considered in the future.

Functionally, a general increase in motor output of ankle extensor muscles could be important for standing and posture as well as for effective weight-bearing during locomotion. While multi-session tSCS did not significantly affect locomotor function as per the BBB scores, due to the nonlinearity of the scale (Fouad et al., 2013; Schucht et al., 2002), an average score of 9.92 for SCI + tSCS animals could still be functionally different compared to the 8.5 for SCI animals. While a score of 8 denotes sweeping motions of the hindlimb and a 9 indicates that there is weight-supported standing on the plantar surface of the paw, a 10 signifies that there is occasional weight-supported plantar stepping (Basso et al., 1995). SCI + tSCS animals had a BBB score leaning toward 10 on average meaning that these animals regained the ability to weight-bear during stance as well as take occasional plantar steps, which is not the case for SCI animals that did not receive tSCS. It appears likely that the increased motor output of the MG muscle contributes to this functional difference, but it will be necessary to incorporate additional measures during stepping such as muscle force, EMG activity, motor patterning, and kinematics to reach that conclusion. More importantly, the tSCS protocol used here was designed to target reflex hyperexcitability after SCI (Knikou, 2013) and was not specifically optimized for improving locomotion and/or rhythmic activity, which appears to require higher stimulation frequency (Gerasimenko et al., 2015a; 2015b; 2015c; Hofstoetter et al., 2015; Minassian et al., 2007b; 2011; 2016a). This is something that will have to be considered moving forward.

Since our knowledge of the effects of tSCS mainly relies on a single stimulation session in SCI individuals (Estes et al., 2018; 2017; Hofstoetter et al., 2014; 2020; Sandler et al., 2021), here, we have investigated the effects of multi-session tSCS in a clinically relevant animal model. For this reason, we wanted to understand how the motor thresholds, recorded in the awake animal, in response to tSCS change over time. Interestingly, we found that motor threshold gradually and significantly decreased over stimulation sessions toward the end of the experimental timeline. It is important to note that this decline in motor threshold in response to a single pulse of tSCS was stable and did not fluctuate erratically from one stimulation session to the next, suggesting that the slight decrease in stimulation threshold that we observed is a true group effect and not a result of variability in electrode placement from animal to animal and from session to session. More importantly, this reduction in motor threshold is combined with decreased H-reflex hyperexcitability, indicating that tSCS does not indiscriminately decrease overall spinal excitability and even aids improvement in motor output. This is further supported by the fact that net motor output is increased following multi-session tSCS after both complete (Malloy et al., 2022) and incomplete SCI (**Figure 6**). However, whether this shift is a result of altered motoneuronal or primary afferent excitability remains to be determined. It would also be interesting to further investigate this change in threshold for activation of motoneurons over time as significant decreases in motor threshold did not occur until 4 weeks post-SCI. Collectively, this supports the use of tSCS acutely after SCI to increase the excitability of motoneuronal activation, which will be beneficial for restoring volitional control and command from spared descending pathways (Gerasimenko et al., 2015a; 2015c; Hofstoetter et al., 2015; Murray et al., 2019; Sayenko et al., 2019; Taccola et al., 2018)

The increased motor output and excitability of motoneuronal activation highlights the major differential improvement of this therapeutic intervention compared to current anti-spastic medications that act to depress central neuronal excitability, including muscle activity (Angeli et al., 2012; Dario and Tomei, 2004; Dietz et al., 2023; Elovic, 2001; Kirshblum, 1999). However, a crux of spasticity after SCI is the impairment of spinal inhibition, thereby rendering reflex pathways and afferent transmission unchecked and exacerbated, so interventions need to also target spinal reflex inhibition. We found that tSCS after injury could decrease hyperreflexia and increase rate modulation of the plantar monosynaptic reflex pathway. Collectively, this indicates that this therapeutic intervention can be used to restore spinal reflex inhibition after SCI without sacrificing a reduction of motor excitability like current anti-spastic medications since, instead, tSCS augments motor output.

#### Nociception is not increased with multi-session tSCS

Maladaptive plasticity within the spinal cord, like central sensitization of nociceptive pathways, can be detrimental to functional recovery and exacerbates the development of hyperexcitability and spasticity (Andresen et al., 2016, Fauss et al., 2022, Finnerup, 2017). It was, therefore, critical to determine whether tSCS induced maladaptive changes in sensory-induced motor responses over time to confirm that the positive effect of tSCS did not come at the cost of hypersensitivity or an increase in nociceptive responses to sensory stimuli. This is especially critical considering the current understanding that tSCS functions by activating dorsal column and dorsal root structures, including cutaneous afferents (Hofstoetter et al., 2018; Holsheimer, 2002). Our results suggest that tSCS does not induce hypersensitivity to tactile and thermal sensory stimuli as the paw withdrawal threshold remained unchanged over time. Additionally, animals did not display any overt nocifensive supraspinal responses and did not show visual signs of pain or discomfort like vocalization, wincing, orbital tightening, or ear folding during stimulation sessions (Sotocinal et al., 2011). On the other hand, spinal cord stimulation has historically been used to address chronic pain with success (Caylor et al., 2019; Illis et al., 1976), suggesting a potential for improvement after SCI. While we did not observe decreased hypersensitivity in the tSCS group, our animal model of severe thoracic contusion is not the most suitable to investigate interventions targeting chronic neuropathic pain and behavioral hypersensitivity after SCI (Eaton, 2003; Verma et al., 2019). Whether this protocol of tSCS after SCI can improve pain, therefore, remains to be determined and will be investigated in a different study.

## CONCLUSION

In summary, tSCS beginning acutely after SCI and lasting 6 weeks post-SCI (1) decreased hyperreflexia and increased plantar H-reflex rate modulation, (2) restored motoneuronal chloride homeostasis, (3) increased motoneuronal gain and motor output, and (4) did not induce hypersensitivity. This work begins to highlight the effects that can be achieved with multiple sessions of tSCS, mechanisms of neuroplastic change, and similarities between tSCS and activity-based therapy, further suggesting it to be a viable intervention for targeting and diminishing the development of spasticity after SCI. Importantly, tSCS differentiates itself from leading pharmacological approaches for spasticity by simultaneously decreasing hyperreflexia and augmenting motor output. The use of this rat model for multi-session tSCS will continue to expand our understanding of the neuroplasticity and functional recovery for spasticity induced by tSCS at an anatomical level, which will be integral for potentiating this intervention as an anti-spastic therapeutic strategy after SCI.

## METHODS AND MATERIALS

All animal procedures were performed in accordance with the National Institutes of Health (NIH) guidelines for the care and use of laboratory animals, and experimental protocols were approved by Drexel University College of Medicine Institutional Animal Care and Use Committee. Thirty-one adult female Sprague Dawley rats (240-260 g; Charles River Laboratories, Wilmington, MA, USA) were used for all experimental procedures. Animals were housed 2-3 per cage with ad libitum food and water under 12-hour light/dark cycle in temperature-controlled facilities accredited by the Association for Assessment and Accreditation of Laboratory Animal Care (AAALAC). All animals were given a one-week acclimation period upon arrival before any procedures were performed. Rats were randomly assigned into one of three groups: neurologically intact (Intact; n = 9), SCI with no stimulation (SCI; n = 11), or SCI with multi-session transcutaneous spinal cord stimulation (SCI + tSCS; n = 11). Terminal experiments occurred 6 weeks post-SCI with Intact animals age-matched on the experimental timeline as controls.

### Surgical procedures and postoperative care

Rats (n = 22) underwent an incomplete contusion spinal cord injury at the thoracic level (T9) under aseptic conditions. Gaseous isoflurane in oxygen (1-4%) was used as an anesthetic prior to and throughout the duration of the surgery. A T8 laminectomy was performed without splitting the dura mater, and stabilization forceps were placed on the spinous processes of the T7 and T9 vertebrae to the Infinite Horizons Impact Device (Precision Systems and Instrumentation, Fairfax, VA, USA). The spinal cord was injured at midline with an average force of 284.44 ± 6.82 kilodynes and a 0 second dwell time by releasing a custom probe (2.0 mm diameter tip) from above the spinal cord. Back muscles and skin incision were sutured accordingly using appropriately sized materials. Rats were singly housed for 3 days following surgical procedures before being paired for the remainder of the study. Animals received one dose of slow-release buprenorphine (0.05 mg/kg, s.c.) as an analgesic prior to surgery end and received saline (5 mL, s.c.) and Baytril (100 mg/kg, s.c.) postoperatively for 7 days to prevent dehydration and infection. Bladders were manually expressed at least twice a day for the duration of the study.

### Behavioral assessments

Animals were acclimated to the individual testing environments and apparatuses for at least one week prior to baseline testing. Locomotor deficits following SCI were assessed via the Basso, Beattie, and Bresnahan (BBB) open-field test using a plastic wading pool (Basso et al., 1995). Each animal was tested individually for 4 minutes, and BBB scores were measured preoperatively (baseline), 3 days post-SCI, 2 weeks post-SCI, 4 weeks post-SCI, and 6 weeks post-SCI. BBB scores ranged from 0 (no hindlimb movement) to 21 (normal movement), and both the left and right hindlimb is scored independently.

Hypersensitivity to tactile and heat stimuli were assessed using dorsal von Frey hair for mechanical allodynia (Detloff et al., 2012) and Hargreaves for thermal hyperalgesia (Hargreaves et al., 1988) tests, respectively. For the dorsal von Frey hair (DVFH) test, animals are securely wrapped in a surgical towel with the hindlimbs exposed. Tactile stimulus is applied perpendicularly to the dorsal surface of the hind paw corresponding to the L5 dermatome using monofilament hairs in ascending size/order from 4.56 (4 g force) to 6.65 (300 g force). Each monofilament is applied 3 times with adequate rest between each application. A positive response is characterized by a rapid withdrawal of the paw in at least two of the three trial applications. This is performed on both left and right hind paws for each animal. DFVH is independent of trunk or hindlimb control (Detloff et al., 2012); therefore, each animal was tested at baseline, 1 week, 2 weeks, 4 weeks, and 6 weeks post-SCI. For the Hargreaves test, animals are placed in a Plexiglas enclosure with mobile heat-flux infrared radiometer machinery located underneath (The Plantar Test Instrument, Ugo Basile, Gemonio, ITA). A noxious heat stimulus is applied to the plantar surface of the hind paw, and withdrawal latency time is recorded automatically upon removal of the limb from above the thermal surface. Each paw (left and right) is tested 5 times with ample rest between each trial. Withdrawal latency times are expressed in seconds, and each reported score reflects an average of three trials per paw after minimum and maximum values were excluded. Hargreaves testing requires weight-bearing of the hind paws on the plantar surface (Hargreaves et al., 1988); therefore, each animal was tested at baseline and any subsequent week where weight-bearing activity in stance on the plantar surface of both hind paws was present.

### Multi-session tSCS

Starting 5 days post-SCI, animals from the SCI + tSCS group were fitted with transcutaneous electrodes as previously described and were secured in a modified, custom-built apparatus to allow them to lie prone with hindlimbs hanging below at rest (Malloy et al., 2022). Briefly, re-usable, self-adhering hydrogel electrodes (Uni-Patch StarBurst Square, Balego, St. Paul, MN, USA) were cut down to 4 cm x 1 cm rectangles, placed over the lumbar enlargement, and covered with 3M Tegaderm transparent film (3M, St. Paul, MN, USA). One electrode (cathode) was placed over the T10-L2 thoracolumbar vertebrae equally between paravertebral sides and no lower than the tips of the T13 floating ribs. Two of the same electrodes, connected to function as a single electrode (anode), were placed bilaterally over the abdomen. The body of the animal was then lightly swaddled with self-adhesive athletic wrap (3M, St. Paul, MN, USA). The motor threshold (T) was evaluated visually and with light manual touch and was determined as the lowest intensity eliciting a twitch of the ankle. Stimulation threshold in milliampere (mA) was recorded for each stimulation session. Stimulations were evoked using a DS3 constant current isolated stimulator (Digitimer, Hertfordshire, GBR) and a customized script written in Signal (CED; Cambridge Electronics Design., Cambridge, GBR). The stimulation protocol consisted of single, monophasic square pulses of 1 ms in duration delivered at 0.2 Hz. Stimulation intensity alternated in bouts of 3 minutes between suprathreshold (1.2 T) and subthreshold (0.8 T) for a total duration of 18 minutes per session. Unstimulated SCI animals were similarly secured to the apparatus for an equal amount of time but were not stimulated. tSCS sessions were repeated 3 times a week for 6 weeks before the terminal experiment.

### Terminal experiments and recordings

Rats were anesthetized with gaseous isoflurane in oxygen (1-4%) and fitted with transcutaneous electrodes as described above. Electromyographic (EMG) needle electrodes (Neuroline Subdermal, Ambu A/S, Ballerup, DNK) were placed in the medal gastrocnemius (MG) and tibialis anterior (TA) muscles, bipolar wire electrodes (Cooner Wire, Chatsworth, CA, USA) were placed in the plantar muscles of the foot on the plantar surface of the hind paw, and ground electrodes were inserted into the skin of the arm. Rats were held at approximately 1.5% isoflurane in oxygen for the duration of the experiment. After completing terminal experiments, rats were sacrificed with an over-dose of Euthasol (150 mg/kg, i.p.).

#### H-reflex

To determine the input-output relationship and assess hyperreflexia, terminal protocols began with a recruitment curve of the H-reflex by recording EMG responses evoked by tibial nerve stimulation. Increasing stimulus intensities allowed for the determination of M-wave and H-wave thresholds, latencies, and maximum response amplitudes recorded from the plantar muscles in the plantar surface of the hind paw. H-reflexes were evoked using an isolated pulse stimulator (Model 2100, A-M Systems, Sequim, WA, USA) delivering single, bipolar pulses of 100 µs duration directly to the tibial nerve. EMG recordings were amplified (x1000; Model 1700, A-M Systems) and band-pass filtered (1 Hz-10 kHz). All signals were digitized (10 kHz) and fed to Signal software (CED).

H-reflex amplitude was kept between 1.2-1.4 times motor threshold (T) and within the ascending potion of the M-wave curve throughout the entire experiment. An intensity of 1.2-1.4 T was used to keep stimulation below the activation threshold for group Ib to group II afferents. Frequency dependent depression (FDD) of the H-reflex was conducted to determine rate modulation capabilities of the monosynaptic reflex pathway. FDD was estimated using a series of 20 consecutive stimulations of the tibial nerve at each increasing frequency of 0.3, 5, and 10 Hz.

#### Transspinal Evoked Potentials (TEPs)

To determine the input-output relationships, terminal protocols began with a recruitment curve of TEPs by recording EMG responses evoked tSCS. Increasing stimulus intensities allowed for the determination of response thresholds, latencies, and maximum response amplitudes recorded from the individual MG and TA hindlimb muscles. TEPs were elicited using the DS7A constant current isolated stimulator (Digitimer, Hertfordshire, GBR) with an analog-to-digital acquisition system and a customized script written in Signal 6 software (CED). Single, monophasic square pulses of 1 ms duration were delivered through the transcutaneous cathodal electrode to the spinal cord. EMG recordings were amplified (x1000; Model 1700, A-M Systems) and band-pass filtered (1 Hz-10 kHz). All signals were digitized (10 kHz) and fed to Signal software (CED). The plantar muscle threshold was used as reference to keep tSCS at 1.2-1.4 T throughout the duration of the experiment.

### Tissue preparation and immunohistochemistry

Rats were perfused transcardially with 0.9% saline followed by 4% paraformaldehyde. Spinal cord was dissected out and stored in the same fixative overnight at 4° C and then incubated in 30% sucrose in PBS for cryoprotection. Tissue was embedded in sectioning medium in preparation for cutting lumbar spinal cord into 30 µm transverse sections on a cryostat. Free-floating sections were blocked in 5% donkey serum and 0.1% Triton X-100 in PBS for one hour. After blocking, sections were incubated overnight at 4° C in the following primary antibodies diluted in 5% donkey serum and 0.1% Triton X-100 in PBS: rabbit anti-potassium-chloride cotransporter 2 (KCC2, 1:400, AB07432, MilliporeSigma, St. Louis, MO, USA) and goat anti-choline acetyl transferase (ChAT, 1:100, AB144P, MilliporeSigma). Sections were washed in PBS and incubated for 2 hours at room temperature in appropriate species-specific donkey secondary antibodies conjugated to Alexa Fluor 488 (1:400, Invitrogen, A21206, Waltham, MA, USA) and Alexa Fluor 647 (1:400, Jackson ImmunoResearch, 705-605-003, West Grove, PA, USA), respectively. Following a final wash, tissue was mounted, cover slipped with Fluoromount-G (SoutherBiotech, Birmingham, AL, USA) and images acquired using a Leica DM550B epifluorescent microscope (Leica Microsystems, Wetzlar, DEU) equipped with a Regita-SRV digital color camera (Q Imaging, Surrey, CAN) using SlideBook version 6.0 imaging software (Intelligent Imaging Innovations, Denver, CO, USA).

### Data analysis

#### BBB

Functional capability via the BBB open-field locomotor test was scored for the left and right hindlimb for each animal. Scores from both hindlimbs were averaged and plotted over baseline, 3 days post-SCI, 2 weeks post-SCI, 4 weeks post-SCI, and 6 weeks post-SCI timepoints for each animal. Group averages were calculated from the averaged hindlimb score of each animal and compared between groups across timepoints.

#### DVFH

Withdrawal threshold for tactile sensation via the DVFH test was recorded in grams of force for both the left and right hind paw of each animal across baseline, 1-, 2-, 4-, and 6-week post-SCI timepoints. Grams of force values for both hind paws were then averaged and normalized to represent a percentage of the baseline withdraw threshold force. Normalized values were used to create group averages for comparison between groups across timepoints.

#### Hargreaves

Hargreaves withdrawal threshold for thermal sensation was recorded in seconds as the latency time to paw withdrawal for both left and right hind paws. Both hind paw latencies were averaged for each animal, and scores were normalized to represent a percentage of the baseline withdrawal threshold time. Group averages were calculated from the averaged withdrawal latency time and compared between groups at 6 weeks post-SCI since this timepoint was the only instance where the maximum number of animals in each group were considered weight-bearing on both hindlimbs.

#### Transcutaneous Spinal Cord Stimulation Thresholds

Stimulation intensity to reach motor threshold was recorded in mA for the SCI + tSCS animals in each of the 18 stimulation sessions. Threshold intensities were then normalized and represented as a percentage of the stimulation intensity required for the first stimulation session. Normalized threshold values were averaged within the group for each session and were compared to session number one to determine the change or stability of motor thresholds during sessions of tSCS over time.

#### Recruitment Curves

Recruitment curves were plotted by expressing the peak-to-peak amplitude of EMG responses as a function of the stimulus intensity. Peak-to-peak amplitude was measured from the maximum negative peak to the maximum positive peak within the duration of the response regardless of the number of peaks. A Boltzmann sigmoid function was then fitted to the curve to estimate parameters of the function such as maximal amplitude (ɑmax), slope of the function (m), and the stimulation intensity requires to reach 50% of ɑmax at a given stimulus intensity (s).

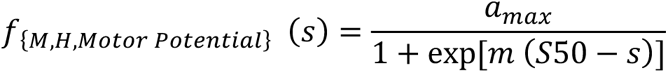

#### H-reflex

M and H response latency (onset of the responses) was calculated by measuring the time between stimulus and response onset. To determine the input-output relationship for the H-reflex, a sigmoid function (Systat SigmaPlot 14.0, Inpixon, Palo Alto, CA, USA) was fitted to the recruitment curve to predict both M-wave and H-wave thresholds, maximal amplitudes (Mmax and Hmax, respectively), slope of the curves, and stimulation intensity required to reach 50% of maximal amplitude. Since Mmax is indicative of the amplitude of response from all plantar muscle motor units innervated by tibial nerve axons and Hmax corresponds to the maximal number of group Ia tibial nerve axons that 28onstitutee the monosynaptic reflex pathway for the plantar muscle (Palmieri et al., 2004), Hmax/Mmax ratios were also measured and compared between groups.

Within the 20 consecutive stimulations of the tibial nerve for FDD of the H-reflex, the first 5 frames were discarded to ensure stabilization of the reflex. Peak-to-peak amplitude was measured as described above and averaged for each animal at 0.3, 5, and 10 Hz. The change in H-reflex amplitude at 5 Hz and 10 Hz was normalized to be calculated as a percentage of the amplitude at 0.3 Hz. Group averages were calculated from the individual normalized values and compared between groups at 5 Hz and 10 Hz.

#### Transspinal Evoked Potentials (TEPs)

To determine the input-output relationship for TEPs, a sigmoid function (Systat SigmaPlot 14.0) was fitted to individual muscle recruitment curves (i.e., each muscle of each animal) to predict stimulation intensity corresponding to threshold, slope, the m function of the slope, maximal amplitude, and stimulation intensity required to reach 50% of maximal amplitude. These parameters were used to normalize individual sigmoid functions to the predicted threshold intensity and maximal amplitude of the curve. Group averages were calculated from the individual normalized values and used to establish a group sigmoid function.

#### KCC2 Membrane Expression

KCC2 immunolabelling was determined by analyzing the fluorescence intensity of the plasma membrane on motoneurons in the ventral horn of the lumbar enlargement. Only large (> 30 µm), ChAT¬+ cells showing a clear nucleus were analyzed. KCC2 fluorescence expression was measured using ImageJ software (National Institutes of Health, Bethesda, MD, USA) by averaging the integrated area under a density curve. Three lines were drawn across the motoneuron from membrane to membrane, each crossing the nucleus, to obtain six total data points per cell (Beverungen et al., 2020; Bilchak et al., 2021; Boulenguez et al., 2010; Côté et al., 2014). The average membrane intensity in each cell was normalized to the cytosolic intensity to determine the amount of KCC2 expressed in the motoneuronal membrane (Boulenguez et al., 2010). Group averages were calculated from the individual normalized ratios and were compared between groups.

### Statistical analysis

Behavioral data such as BBB, DFVH, and Hargreaves were compared between groups and across timepoints using a two-way mixed-design (Mixed) analysis of variance (ANOVA) followed by the Holm-Sidak post hoc multiple comparisons test. Threshold of stimulation intensity for each tSCS session in the SCI + tSCS group was compared to the intensity required for session one using a Friedman one-way repeated measures (RM) ANOVA. Friedman’s test was used since normality or equal variance tests failed. For H-reflex recruitment curves, amplitude was normalized as a percentage of Mmax and plotted as a function of stimulus intensity representing multiples of motor threshold (xMT). For TEP recruitment curves of each muscle, amplitude was normalized as a percentage of maximal amplitude and plotted as a function of stimulus intensity (xMT). A two-way RM ANOVA followed by Holm-Sidak post hoc test was utilized to compare amplitudes at increasing stimulation intensities between Intact, SCI, and SCI + tSCS groups for both the H-reflex and TEPs. A two-way RM ANOVA followed by Holm-Sidak post hoc test was also used to compare H-reflex amplitude during FDD between groups.

Differences in all recruitment curve parameters as well as TEP latency and M-wave and H-wave thresholds, latencies, maximal amplitudes, and Hmax/Mmax ratios between Intact, SCI, and SCI + tSCS groups were determined using a one-way ANOVA followed by Holm-Sidak post hoc test. Kruskal-Wallis one-way ANOVA followed by Dunn’s post hoc multiple comparison test was used if normality or equal variance tests failed. A Kruskal-Wallis one-way ANOVA followed by Dunn’s post hoc test was also used to compare KCC2 membrane expression ratios between groups. Statistical analyses were performed using GraphPad Prism software version 7.04 (GraphPad Software, San Diego, CA, USA). For all tests, significance was determined when p < 0.05, and values are reported as the group average ± standard error of the mean.

## Acknowledgments

This work was supported by grants from the National Institute of Neurological Disorders and Stroke (RO1 NS119475) and the Craig H. Neilsen Foundation (647897). We thank Drs Maria Knikou and Simon Danner for helpful comments on an earlier version of this manuscript.

